# Cytotoxic lymphocytes target HIV-1 Gag through granzyme M-mediated cleavage

**DOI:** 10.1101/2021.02.24.432686

**Authors:** Elisa Saccon, Flora Mikaeloff, Pol Figueras Ivern, Ákos Végvári, Anders Sönnerborg, Ujjwal Neogi, Robert van Domselaar

## Abstract

HIV-1 leads to progression to immunodeficiency and death of individuals who do not receive successful antiretroviral therapy. Initially, the host’s immune response controls the infection, but cannot eliminate the HIV-1 from the host. Cytotoxic lymphocytes are the key effector cells in this response and can mediate crucial antiviral responses through the release of a set of proteases called granzymes towards HIV-1-infected cells. However, little is known about the immunological molecular mechanisms by which granzymes could control HIV-1. Since we noted that HIV-1 subtype C (HIV-1C) Gag with the tetrapeptide insertion PYKE contains a putative granzyme M (GrM) cleavage site (KEPL) that overlaps with the PYKE insertion, we analyzed the proteolytic activity of GrM towards Gag. Immunoblot analysis showed that GrM could cleave Gag proteins from HIV-1B and variants from HIV-1C of which the Gag-PYKE variant was cleaved with extremely high efficiency. The main cleavage site was directly after the insertion after leucine residue 483. GrM-mediated cleavage of Gag was also observed in co-cultures using cytotoxic lymphocytes as effector cells and this cleavage could be inhibited by a GrM inhibitor peptide. Altogether, our data indicate towards a noncytotoxic immunological mechanism by which GrM-positive cytotoxic lymphocytes target the HIV-1 Gag protein within infected cells to potentially control HIV-1 infection. This mechanism could be exploited in new therapeutic strategies to treat HIV-1-infected patients to improve immunological control of the infection.

## Introduction

A key component of both the innate and adaptive immune response against viruses is represented by cell-mediated cytotoxicity, mediated by cytotoxic lymphocytes, through the release of granules containing perforin and granzymes. Granzymes are a family of serine proteases, that can be expressed by cytotoxic lymphocytes, such as natural killer (NK) cells and CD8^+^ T cells. During the early phases of infection, granzymes enter the virus-infected target cells, facilitated by pore-forming protein perforin, and mediate antiviral effects through cleavage of host and/or viral proteins. In humans, five granzymes are encoded, namely GrA, GrB, GrH, GrK, and GrM (1–4). These five granzymes differ in their antiviral functions through their difference in substrate specificity (5). Specifically, GrM cleaves after a leucine or methionine and its four amino acid consensus substrate motif is KEPL (6, 7). GrM can mediate antiviral effects through induction of apoptosis or by inhibiting viral replication independent of cell death (4, 8, 9). Noncytotoxic functions of granzymes have been shown to be of importance in controlling chronic and latent infections, such as herpes simplex virus type I and cytomegalovirus infections (8–10). On the other side, little is known about the role of granzymes in HIV-1 infection. A recent study showed that GrM protein expression in HIV-1 specific CD8^+^ T cells from HIV-1 infected individuals was the highest compared to expression of the other granzymes (11). Furthermore, killing of HIV-1-infected CD4^+^ T cells by HIV-1-specific CD8^+^ T cells was perforin-dependent but GrB-independent because of the expression of GrB-inhibitor SerpinB9 within the target cells. Which other granzymes could be responsible for the killing of the HIV-1-infected CD4^+^ T cells and whether these granzymes employ noncytotoxic antiviral functions remain unclear. These granzyme-mediated antiviral mechanism could be exploited or boosted in HIV-1-infected individuals to provide a better immunological control of their infection.

Among all HIV-1 subtypes, HIV-1 subtype B (HIV-1B) is the most prevalent in most high-income countries. However, subtype C (HIV-1C) causes over 50% of all HIV-1 infections worldwide and has become increasingly prevalent in Europe (12, 13). Recently, a tetrapeptide insertion PYKE within a relatively conserved region of the viral Gag protein (Gag_PYKEi_) was identified in a subset of HIV-1C-infected individuals (14). HIV-1 protein Gag mediates the assembly, budding, and maturation of new virions (15). The PYKE motif is important for the interaction of the viral Gag to host cell protein ALIX that facilitates viral budding. Furthermore, insertion of this motif correlated with enhanced viral fitness (16, 17). Interestingly, HIV-1C Gag_PYKEi_ contains the GrM consensus substrate motif KEPL overlapping the PYKE motif, whereas HIV-1B and HIV-1C without the tetrapeptide insertion only contain the proline and leucine residues at this site within Gag. Thus, HIV-1C Gag_PYKEi_ and perhaps other Gag variants are putative GrM substrates, and cleavage by GrM could then potentially interfere with budding and infectiousness of new HIV-1 virions.

In this study, we further investigated the role of GrM in HIV-1 infection. First, we evaluated the proteolytic potential of GrM to target the Gag protein from various HIV-1 subtypes, including HIV-1C with or without the tetrapeptide insertion. Then, we assessed whether GrM can target the Gag protein in an *in vitro* cell model. Finally, we examined the gene and protein expression of GrM within PBMCs of a cohort of HIV-1 infected individuals. Specifically, we compared the proteo-transcriptomic profiles of Elite controllers (ECs) to viral progressors (VPs) and uninfected individuals (HC), to assess whether differences in GrM levels could be an underlying immunological mechanism by which ECs can control their HIV-1 infection in the absence of antiretroviral therapy.

## Material and Methods

### Cell culture

Cells were cultured in 5% CO_2_ at 37°C. HEK293FT cells were maintained in Dulbecco’s modified Eagle medium (DMEM, Gibco/ThermoFisher Scientific, USA) supplemented with 10% fetal bovine serum (FBS, Sigma, USA), 2 mM L-glutamine (Sigma, USA), 0.1 mM MEM Non-Essential Amino Acids (Gibco/ThermoFisher Scientific, USA), and 20 units/mL penicillin combined with 20 μg/mL streptomycin (Sigma, USA). HeLa cells (#ATCC CCL-2) were maintained in DMEM supplemented with 10% FBS and 20 units/mL penicillin combined with 20 μg/mL streptomycin. KHYG-1 cells (#ACC 725, DSMZ, Germany) were maintained in Roswell Park Memorial Institute 1640 (RPMI, Sigma, USA) medium supplemented with 10% FBS, 25 mM HEPES, 20 units/mL penicillin and 20 μg/mL streptomycin, and 100 units/mL of recombinant human interleukin-2 (IL-2, PeptroTech, Sweden).

### Plasmids

pCR3.1/HIV1B-Gag-mCherry (HIV-1B Gag) was a kind gift from Dr. Paul Bieniasz (The Rockefeller University, USA). pCR3.1/HIV1C-Gag-mCherry variants with or without PYKE or PYQE tetrapeptide insertions were described previously (16). The Gag_PYKEi-L483A_ mutant was generated by PCR-directed cloning using pCR3.1/HIV-GagPYKEi-mCherry as template. All other Gag_PYKEi_ mutants were generated by PCR-directed cloning using the L483A mutant as template. PCR was performed using Phusion DNA polymerase (NEB) and corresponding primers (Table I). Linear amplicons were circularized using T4 polynucleotide kinase and T4 ligase. DNA sequences from all constructs were verified by Sanger sequencing.

**Table I.**
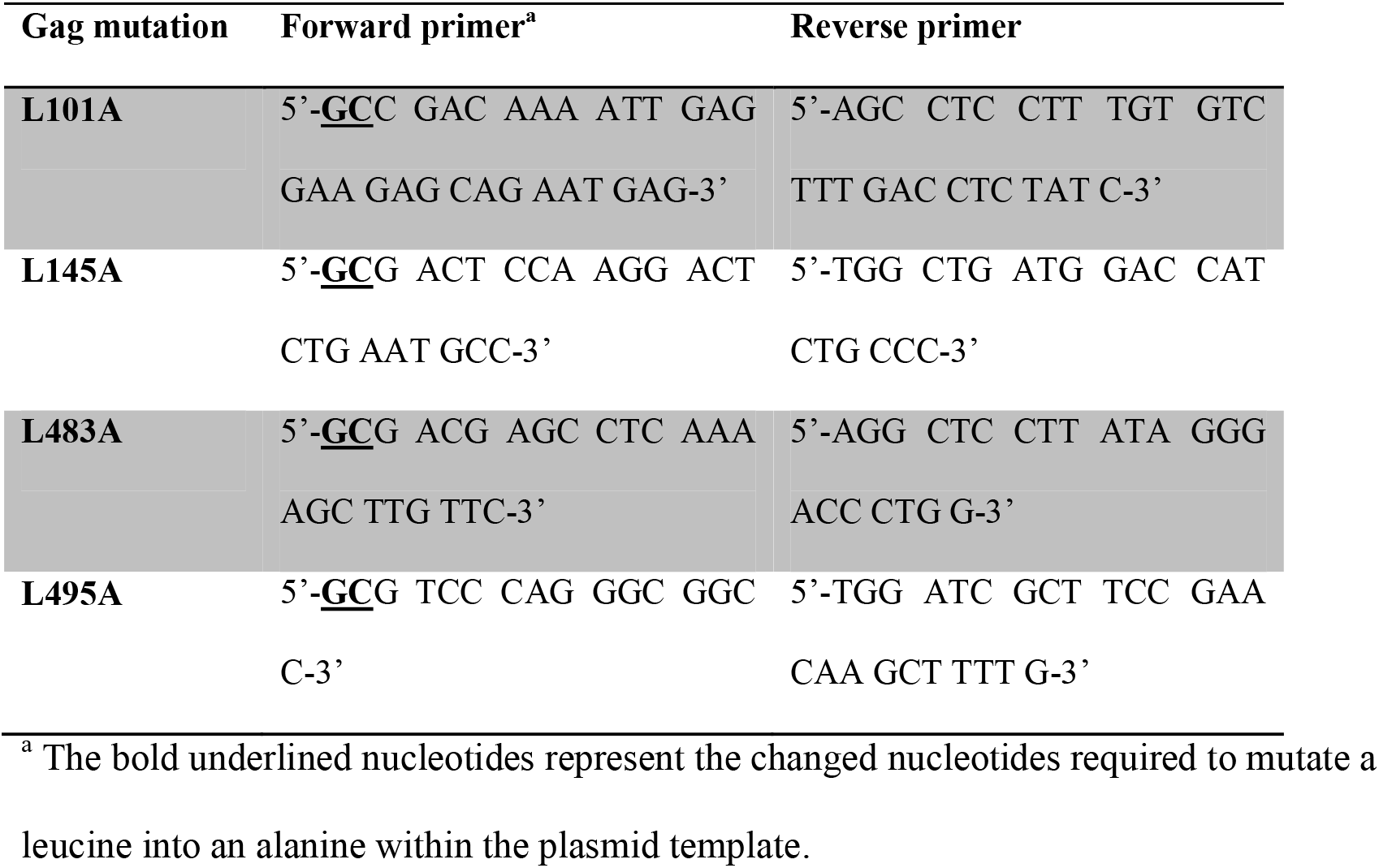
Primers used for mutagenesis of Gag.

### Antibodies and reagents

Rabbit monoclonal antibody against GFP (EPR14104, ab183734), rabbit polyclonal antibody against mCherry (ab167453), and rabbit polyclonal to HIV-1 gag (p55 + p24 + p17, ab63917) were purchased from Abcam (UK). Horse-radish peroxidase-conjugated rabbit anti-goat was obtained from Agilent Dako (USA). NuPAGE™ 4-12% Bis-Tris Protein Gels and iBlot™ Transfer Stacks were purchased from ThermoFisher Scientific (USA). Immunoblotted proteins were detected using the Enhanced Chemiluminescence detection system (Amersham, UK) and ChemiDoc XRS+ (Bio-Rad, USA). Pan-caspase inhibitor zVAD-FMK was purchased from Enzo Life Sciences (USA) and GrM inhibitor AcKVPL-CMK from Peptanova (Germany).

### GrM cleavage assay

Purified recombinant catalytically active human GrM and the catalytically inactive GrM variant (GrM-SA) were a kind gift from Dr. N. Bovenschen (University Medical Center Utrecht, The Netherlands). HEK293FT cells (4×10^6^) were seeded into a 10cm cell culture dish one day prior to transfection. Then, cells were transfected with one of the Gag-mCherry variants (8 µg) using FuGene HD according to the manufacturer’s instructions (Promega, USA) in a 3:1 ratio with DNA. One day post-transfection, cell-free protein lysates were generated by washing cells three times in PBS and subsequent lysis in PBS by three cycles of freeze-thawing using liquid nitrogen. Samples were centrifuged at 18,000 x *g* for 10 min at 4°C, supernatant was collected, aliquoted and stored at −20°C. Protein concentration was determined by DC protein assay (Bio-Rad, USA). Lysates (10 µg) were incubated with indicated concentrations of either GrM or GrM-SA supplemented with a Tris-buffer (50 mM Tris-HCl pH 7.4 and 150 mM NaCl) to a total volume of 12 µL for 4 hours (or otherwise indicated) at 37°C. Then, 4x Leammli buffer was added, samples were incubated at 95°C for 10 min and subjected to immunoblotting. ImageLab software (Bio-Rad, USA) was used to quantify protein band intensities and GraphPad Prism to plot values in a graph.

### KHYG-1 and IL-2-activated PBMC killer cell assay

PBMCs were isolated from buffy coats of anonymous donors using Ficoll density centrifugation, aliquoted and stored in liquid nitrogen. A frozen aliquot of PBMCs was thawed and incubated for 4 days in RPMI 1640 medium (Gibco, USA) supplemented with 5% human AB serum (Sigma, USA) and 1000 units/mL of recombinant human IL-2. HeLa cells (2×10^6^) were collected and transfected with pCR3.1/HIV-GagPYKEi-mCherry (7.5 µg) and pEGFP-N1 (2.5 µg) and directly seeded into a 96-wells plate (20,000 cells per well). When the pan-caspase inhibitor zVAD-FMK was used, seeded cells were incubated with 100 µM of zVAD-FMK or only DMSO. One day post-transfection, transfected HeLa cells were challenged with either KHYG-1 cells or IL-2-activated PBMCs in indicated effector:target (E:T) ratios for in 5% CO_2_ at 37°C. When GrM inhibitor AcKVPL-CMK was used, KHYG-1 were pre-treated with 100 µM of AcKVPL-CMK for 1 hour, before the co-culture with target cells was started. After indicated times, cells were washed twice with PBS and directly lysed in 2x Laemmli buffer. Samples were incubated at 95°C for 10 min. and subjected to immunoblotting. ImageLab software was used to quantify protein band intensities and GraphPad Prism was used to plot values in a graph.

### Gene expression analysis on patient samples

Whole blood was obtained from HIV-1-negative donors (HC, n=16), untreated HIV-1-infected individuals without viremia (Elite Controllers, EC, n=19), and treatment naïve HIV-1-infected patients with viremia (Viremic Progressors, VP, n=17) as described before (18). The criteria to be classified as EC is that the patient has been diagnosed as HIV-1 seropositive for over a year with ≥3 consecutive viral loads of <75 viral RNA copies/mL (with all previous viral loads below 1000 copies/mL) or has been HIV-1 seropositive for over 10 years with a minimum of two viral loads (and in total 90% of all viral loads) of 400 copies/mL. In case of the ECs, the viral load was below detection limit (<20 copies/mL) at time of sampling. Peripheral blood mononuclear cells (PBMCs) were isolated from whole blood through Ficoll density centrifugation (Ficoll-Plaque PLUS, GE Healthcare, USA), aliquoted and stored in liquid nitrogen. RNA sequencing on extracted RNA from PBMCs was performed as described previously (18). Then, data was extracted using Ensembl transcript IDs (Table II) and normalized transcript level values for each patient within each patient group were plotted in graphs.

**Table II.** Transcript and protein IDs used for extracting data regarding transcript and protein levels for all human granzymes and perforin.

| Protein name | Gene name | Transcript ID | bp | UniProt | aa |
| --- | --- | --- | --- | --- | --- |
| **Granzyme A** | *GZMA* | ENST00000274306.7 (GZMA-201) | 896 | P12544 | 262 |
| **Granzyme B** | *GZMB* | ENST00000216341.9 (GZMB-201) | 891 | P10144 | 247 |
| **Granzyme H** | *GZMH* | ENST00000216338.9 (GZMH-201) | 936 | P20718 | 246 |
| **Granzyme K** | *GZMK* | ENST00000231009.2 (GZMK-201) | 1509 | P49863 | 264 |
| **Granzyme M** | *GZMM* | ENST00000264553.6 (GZMM-201) | 924 | P51124 | 257 |
| **Perforin** | *PRF1* | ENST00000373209.2 (PRF1-201) | 2492 | P14222 | 555 |

bp, base pairs; aa, amino acids.

### Protein expression analysis on patient samples using LC-MS/MS

Proteomic analysis was performed on extracted proteins from PBMCs of nine male individuals from each group. Cell pellets were suspended in 40 µL of 0.1% ProteaseMAX™ (Promega, Madison, WI) and 4M urea in 50 mM ammonium bicarbonate and 10% acetonitrile (ACN). The samples were sonicated using Vibra-Cell™ probe (Sonics & Materials, Inc., Newtown, CT) for 1 min, with pulse 2/2, at 20% amplitude, and sonicated in bath for 5 minutes, followed by vortexing and centrifugation for 5 min at 13,000 rpm. The supernatants yielded 3-95 µg proteins.

Ten µg of each sample (except for the samples we designated as EC1 (7 µg), HC13 (3 µg), HC17 (3 µg), VP3 (8 µg) and VP10 (3 µg)) were subjected to proteolytic digestion following reduction in 6 mM dithiothreitol at 37°C for 60 min and alkylation in 22 mM iodoacetamide for 30 min at room temperature in the dark. Trypsin was added as enzyme to a protein ratio of 1:50 and digestion was carried out at 37°C over night. Tryptic peptides were cleaned on Thermo Scientific™ HyperSep™ C18 Filter Plate, bed volume 40 µL (ThermoFisher Scientific, USA) and dried in a centrifugal concentrator (Genevac™ miVac, Fisher Scientific, USA).

Thermo Scientific™ TMT10plex™ isobaric label reagents (ThermoFisher Scientific, USA) in 100 µg aliquots were dissolved in 30 µL dry ACN, scrambled and mixed with the digested samples solubilized in 70 µL triethylammonium bicarbonate, followed by incubation at 22°C for 2 h at 550 rpm. The reaction was quenched with 12 µL of 5% hydroxylamine at 22°C for 15 min at 550 rpm. The labeled samples were pooled and dried in a centrifugal concentrator. The TMT-labeled tryptic peptides were dissolved in 20 µL of 2% ACN/0.1% formic acid. Reversed-phase liquid chromatography was performed on a Thermo Scientific™ EASY-nLC 1000 system (ThermoFisher Scientific, USA) on-line coupled to a Q Exactive™ Plus Hybrid Quadrupole-Orbitrap™ mass spectrometer (ThermoFisher Scientific, Bremen, Germany). Samples were injected onto a 50 cm long C18 Thermo Scientific™ EASY-Spray™ column (ThermoFisher Scientific, USA) and separated with the following gradient: 4-26% ACN in 180 min, 26-95% ACN in 5 min, and 95% ACN for 8 min at 300 nL/min flow rate. Mass spectrometric data acquisition was comprised of one survey mass spectrum in *m/z* 350 to 1600 range, acquired with 140,000 (at *m/z* 200) resolution, followed by higher energy collision dissociation (HCD) fragmentations of the 16 most intense precursor ions with 2+ and 3+ charge state, using 60 s dynamic exclusion. The tandem mass spectra were acquired with 70,000 resolution, targeting 2×10^5^ ions, using *m/z* 2.0 isolation width and 33% normalized collision energy.

The raw data files were directly loaded in Thermo Scientific™ Proteome Discoverer™ v2.2 (ThermoFisher Scientific, San José, CA) and searched against human SwissProt protein databases (21,008 entries) using the Mascot Server 2.5.1 search engine (Matrix Science Ltd., London, UK). Search parameters were chosen as follows: up to two missed cleavage sites for trypsin, mass tolerance of precursor and HCD fragment ions at 10 ppm and 0.05 Da, respectively. Dynamic modifications of oxidation on methionine, deamidation of asparagine and glutamine and acetylation of N-termini were set. For quantification, both unique and razor peptides were requested. Protein raw data abundance was first filtered with an in-house script and quantile normalized with NormalizerDE v1.4.0 (19). Histogram was used to assess that the distribution follows a normal law. The batch effect of multiple TMT experiments was removed using the ComBat function of the sva R package v3.34.0 (20). Differential expression analysis was performed with R package limma v3.42.2 to determine protein abundances (21). Benjamini-Hochberg (BH) adjustment and 0.05 FDR cut-off was applied. Finally, data was extracted using the Uniprot IDs (Table II) and normalized protein abundances for each patient within each patient group were plotted in graphs.

### Ethics statement

The study was approved by regional ethics committees of Stockholm (2013/1944–31/4) and amendment (2019-05585). All participants gave written informed consent. Patient identity was anonymized and de-identified before analysis.

## Results

### GrM targets various HIV-1 Gag variants

As HIV-1C Gag_PYKEi_ contains the GrM consensus substrate motif KEPL, we first assessed whether Gag_PYKEi_ could be cleaved by GrM. Since the predicted cleavage site is within the P6 late domain of Gag, we used an HIV-1C Gag_PYKEi_ cDNA construct with mCherry fused to its C-terminus. We prepared cell-free protein lysates with overexpressed mCherry-tagged Gag_PYKEi_, incubated them with GrM and assessed GrM-mediated Gag_PYKEi_ cleavage by immunoblotting using antibodies against either mCherry or Gag. Whereas the catalytic inactive GrM control (GrM-SA) did not cleave Gag_PYKEi_, the viral protein was cleaved in a dose-(Figure 1A-B) and time-dependent manner (Figure 1C-D) by GrM. Already at a very low concentration of 5 nM, GrM almost completely degraded full-length Gag_PYKEi_ within one hour. Immunoblotting using an mCherry antibody showed a degradation product of around 29 kDa, which could represent the product of mCherry with a small part of the P6 late domain. Immunoblotting using a polyclonal antibody that detects the matrix and capsid domains of Gag revealed degradation products of around 53 kDa, which could represent the Gag_PYKEi_ without the last part of the P6 late domain, and 40 kDa. This indicates that HIV-1C Gag_PYKEi_ can be targeted by GrM. Moreover, the presence of multiple cleavage products suggests that GrM proteolytically processed Gag_PYKEi_ at least at two different sites.

**Figure 1.**
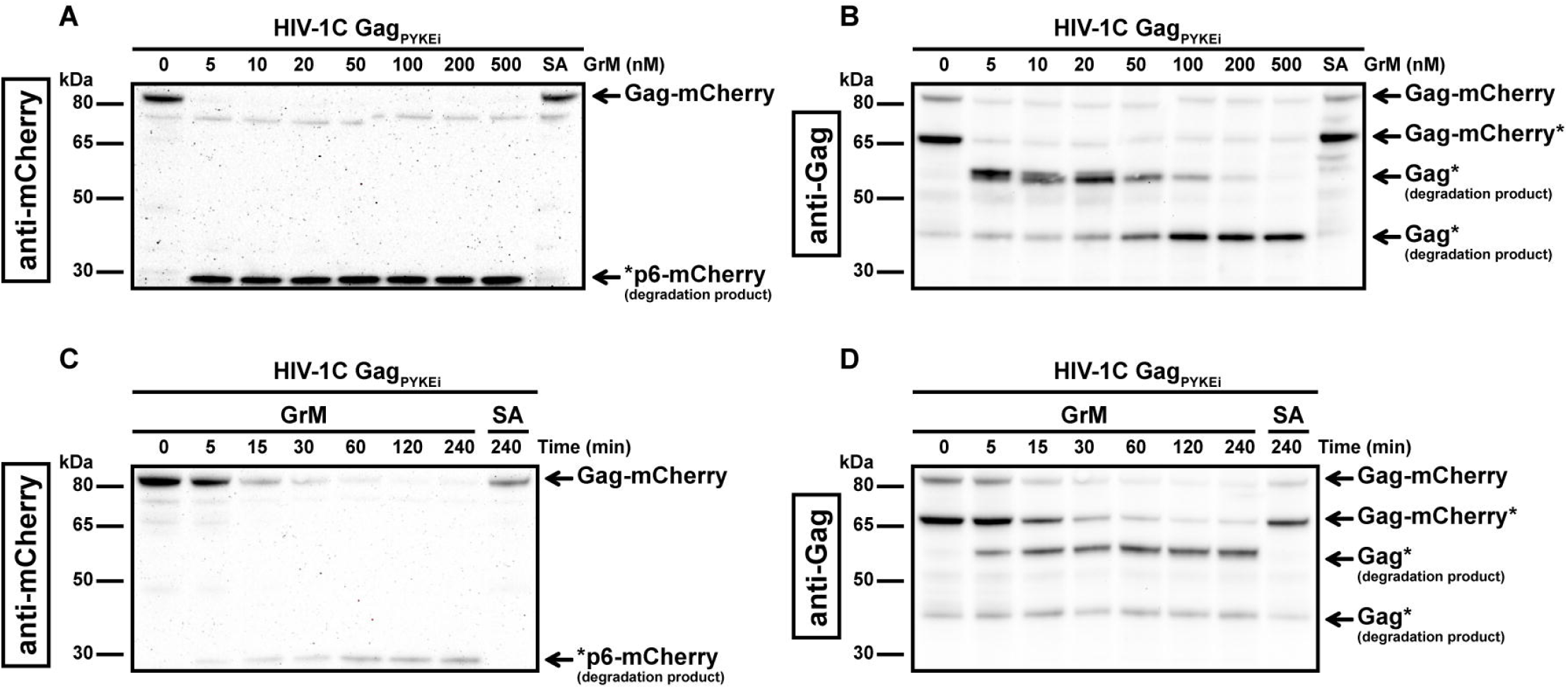
GrM degrades HIV-1 Gag_PYKEi_ protein. **(A-B)** HEK293FT cells were transfected with plasmids encoding for C-terminal mCherry-tagged HIV-1C Gag_PYKEi_ and lysates (10 µg) were incubated with increasing concentrations of GrM or GrM-SA (500 nM) for 4 h at 37°C. Samples were subjected to immunoblotting using an anti-mCherry antibody **(A)** or anti-p55 Gag antibody **(B)** to detect full length Gag-mCherry and degradation products. **(C-D)** Lysates (10 µg) were incubated with 5 nM of GrM for the indicated time points or GrM-SA for 4 h at 37°C and immunoblotted using an anti-mCherry antibody **(C)** or anti-p55 Gag antibody **(D)**. Of note, the C-terminal mCherry tag is partially degraded by other cellular proteases as observed by the smaller Gag-mCherry product around 65 kDa (Gag-mCherry*). Data depicted is representable for at least two individual experiments.

Next, we examined the ability of GrM to cleave other Gag variants. When we tested GrM-mediated cleavage of Gag variants HIV-1C with a PYQE insertion (HIV-1C Gag_PYQEi_) (Figure 2A-B) or without any tetrapeptide insertion (HIV-1C Gag_wt_) (Figure 2C-D) and HIV-1B (Figure 2E-F), we observed degradation of Gag with GrM concentrations of 20 nM or higher. Cleavage is most efficient in the HIV-1C Gag_PYKEi_ variant, which contains the complete consensus GrM substrate motif (Figure 2G). This is followed by HIV-1C Gag_PYKEi_, HIV-1B and finally HIV-1C wild type in terms of proteolytic efficiency by which GrM can degrade the Gag protein. In conclusion, these data indicate that GrM can cleave Gag regardless of subtype.

**Figure 2.**
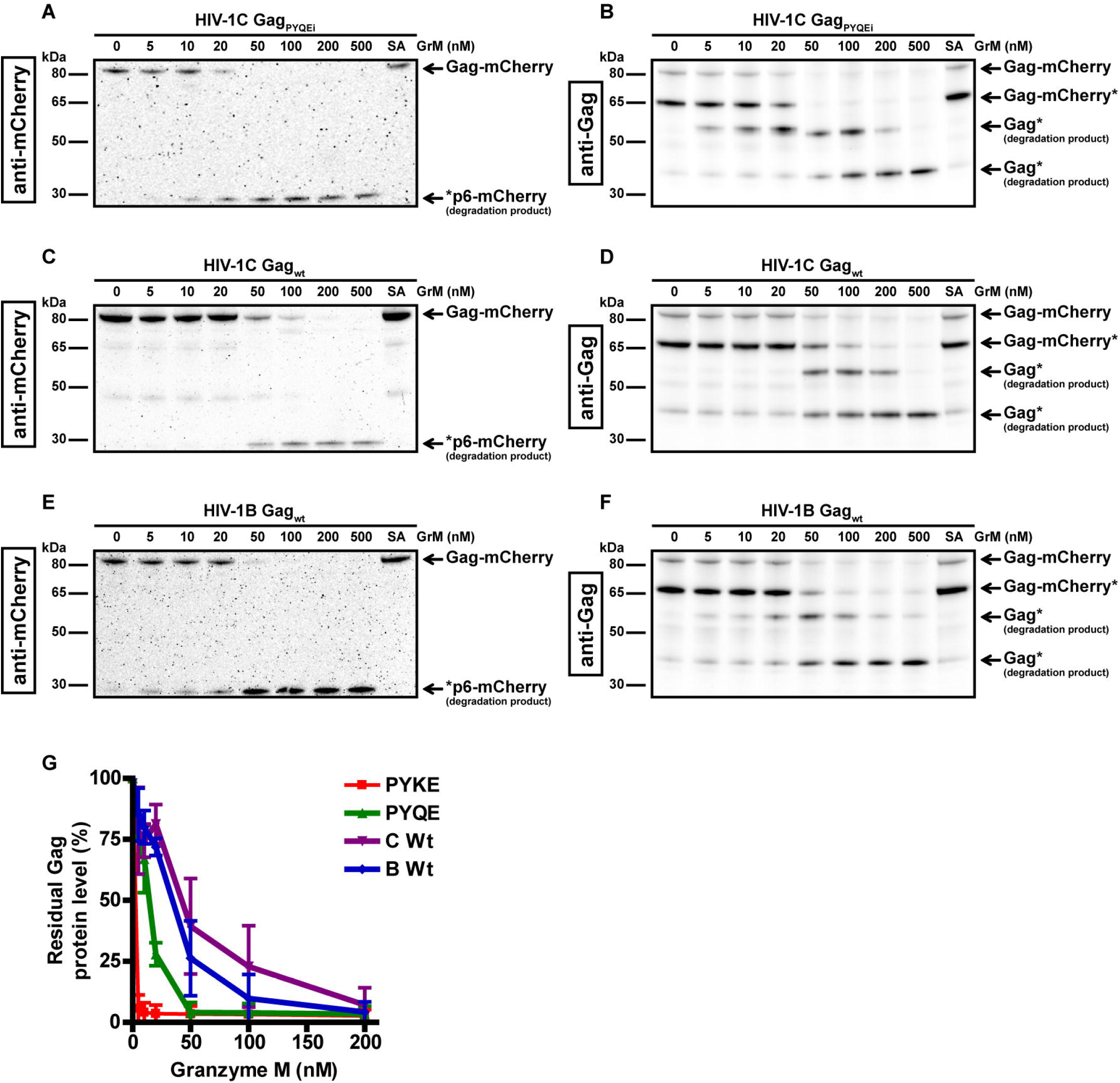
GrM degrades various HIV-1 subtype Gag proteins. HEK293FT cells were transfected with plasmids encoding for C-terminal mCherry-tagged HIV-1C Gag_PYQEi_ **(A-B)**, HIV-1C Gag_WT_ **(C-D)** or HIV-1B Gag_WT_ **(E-F)** and lysates (10 µg) were incubated with increasing concentrations of GrM or GrM-SA (500 nM) for 4 h at 37°C. Samples were subjected to immunoblotting using an anti-mCherry antibody **(A, C, E)** or anti-p55 Gag antibody **(B, D, F)** to detect full length Gag-mCherry and degradation products. **(G)** Band intensities of full-length Gag-mCherry from all four Gag variants as detected with the anti-mCherry antibody in figures 1 and 2 and additional experiments were quantified and plotted with Gag incubated with 0 nM GrM set at 100%. Data points represent the mean ±SEM from two individual experiments.

### Tetrapeptide insertion PYKE facilitates very efficient GrM-mediated cleavage after Leu^483^

The difference in cleavage efficiency between the tested HIV-1C variants could be caused by the variation in protein sequence at the site of the tetrapeptide insertion. The PYKE insertion in HIV-1C Gag creates a complete GrM consensus substrate motif KEPL. Therefore, we used this variant as a tool for our next experiments to identify the main GrM cleavage site and to assess GrM-mediated cleavage of Gag in living cells. First, to test whether GrM can indeed cleave Gag after the predicted leucine residue, we mutated this leucine residue within Gag_PYKEi_ (Leu^483^) into an alanine (PYKEi^L483A^). HIV-1C-Gag_PYKEi_ and PYKEi^L483A^ were expressed in HEK293FT cells and cell-free protein extracts were incubated with increasing concentrations of GrM (Figure 3A-B). Whereas Gag_PYKEi_ was very efficiently cleaved by GrM, the cleavage efficiency of the PYKEi^L483A^ mutant was significantly lower.

**Figure 3.**
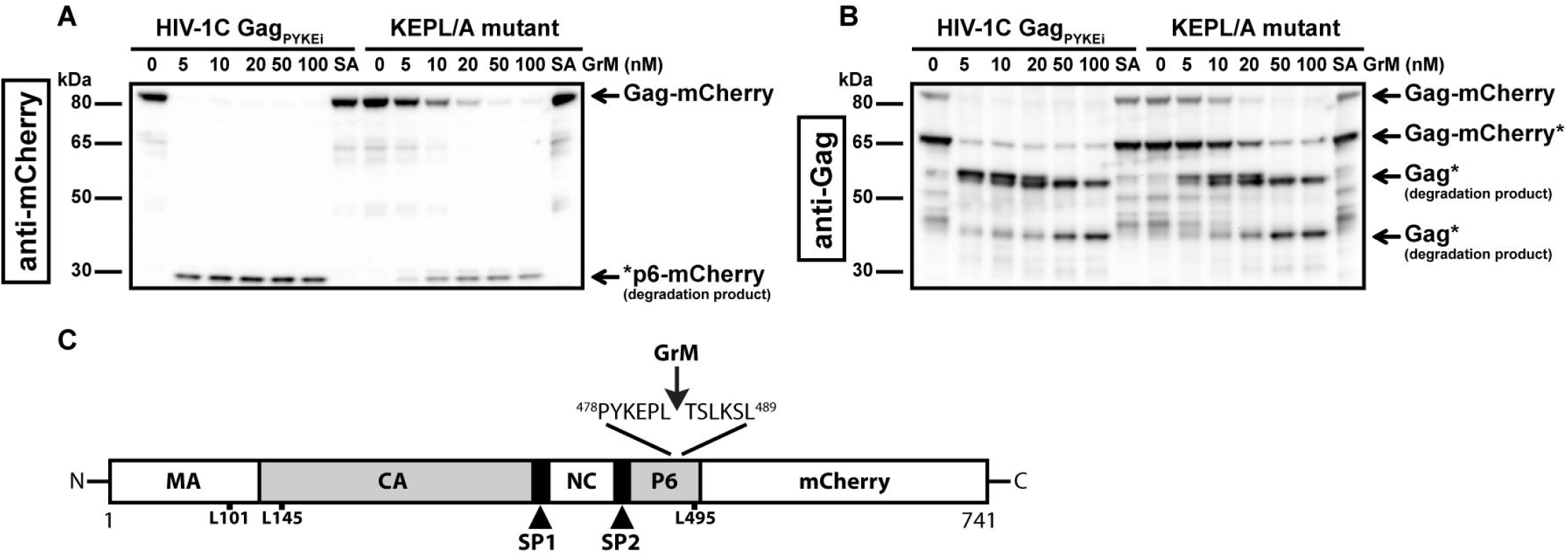
GrM cleaves HIV-1C Gag_PYKEi_ after Leu^483^. **(A-B)** Based on the HIV-1C Gag_PYKEi_ protein sequence and the GrM acid consensus GrM substrate motif KEPL, we predicted that GrM can cleave Gag_PYKEi_ after the leucine residue at position 483 (Leu^483^). To test this, we mutated Leu^483^ into an alanine (KEPL/A mutant). Then, HEK293FT cells were transfected with mCherry-tagged HIV-1C Gag_PYKEi_ or the KEPL/A mutant. Lysates (10 µg) were incubated with increasing concentrations of GrM or GrM-SA (100 nM) for 4 h at 37°C and immunoblotted using an anti-mCherry antibody **(A)** or anti-p55 Gag antibody **(B). (C)** Schematic overview of GrM cleavage site within Gag_PYKEi_ as well as other tested mutants that were not GrM cleavage sites (MA, p17 matrix protein; CA, p24 capsid protein; SP1, spacer peptide 1; NC, p7 nucleocapsid protein; SP2, spacer peptide 2; P6, p6 late domain).

Since we observed that GrM cleaves Gag_PYKEi_ at multiple sites, albeit at lower efficiencies, we also tested whether GrM cleaves at other potential sites. Based on the size of cleavage products, we predicted there is another cleavage adjacent to the identified Leu^483^ residue as well as at the end of the matrix region or beginning of the capsid region. We mutated residues Leu^101^, Leu^145^ or Leu^495^ within PYKEi^L483A^ into an alanine, prepared cell-free protein extracts from cells overexpressing each mutant variant and incubated these extracts with GrM. None of these mutants showed a significant resistance in GrM-mediated cleavage (data not shown). Altogether, Gag_PYKEi_ is most efficiently cleaved by GrM after Leu^483^ (Figure 3C).

### Cytotoxic lymphocytes target Gag_PYKEi_ through GrM in living cells

Our previous experiments to analyze GrM-mediated cleavage of Gag were performed in *in vitro* lysates and purified GrM. Next, we wanted to assess whether GrM could target Gag in a more immunologically relevant setting using co-culture assays with target cells challenged by cytotoxic lymphocytes. To this end, we overexpressed Gag_PYKEi_ and GFP in HeLa cells and co-cultured these target cells with IL-2-activated PBMCs, which express all granzymes. After four hours of co-culture, the supernatant was removed, and the remaining adherent cells were directly lysed and subjected to immunoblotting. Since IL-2-activated PBMCs induce non-specific target cell death, but will still be present in our lysates, we used the GFP expression in the HeLa cells as cell viability loading control. Indeed, with increasing effector:target (E:T) ratios, more cell death was induced as observed by the decrease in GFP levels (Figure 4A). No additional decrease in full-length Gag_PYKEi_ nor appearances of degradation products were observed. To limit the bias of cell death in our assay, we performed the experiment in the presence of pan-caspase inhibitor zVAD-FMK. Co-culture in the presence of this pan-caspase inhibitor inhibited cell death (Figure 4B). Although there was no clear decrease in full length Gag, we did observe a degradation product of around ~53 kDa similar to the *in vitro* lysate experiments. Of note, *de novo* Gag is transported to the membrane where it forms vesicles and is thus constantly released from cells (22), and this could explain why a decrease of full-length Gag through degradation is not observed.

**Figure 4.**
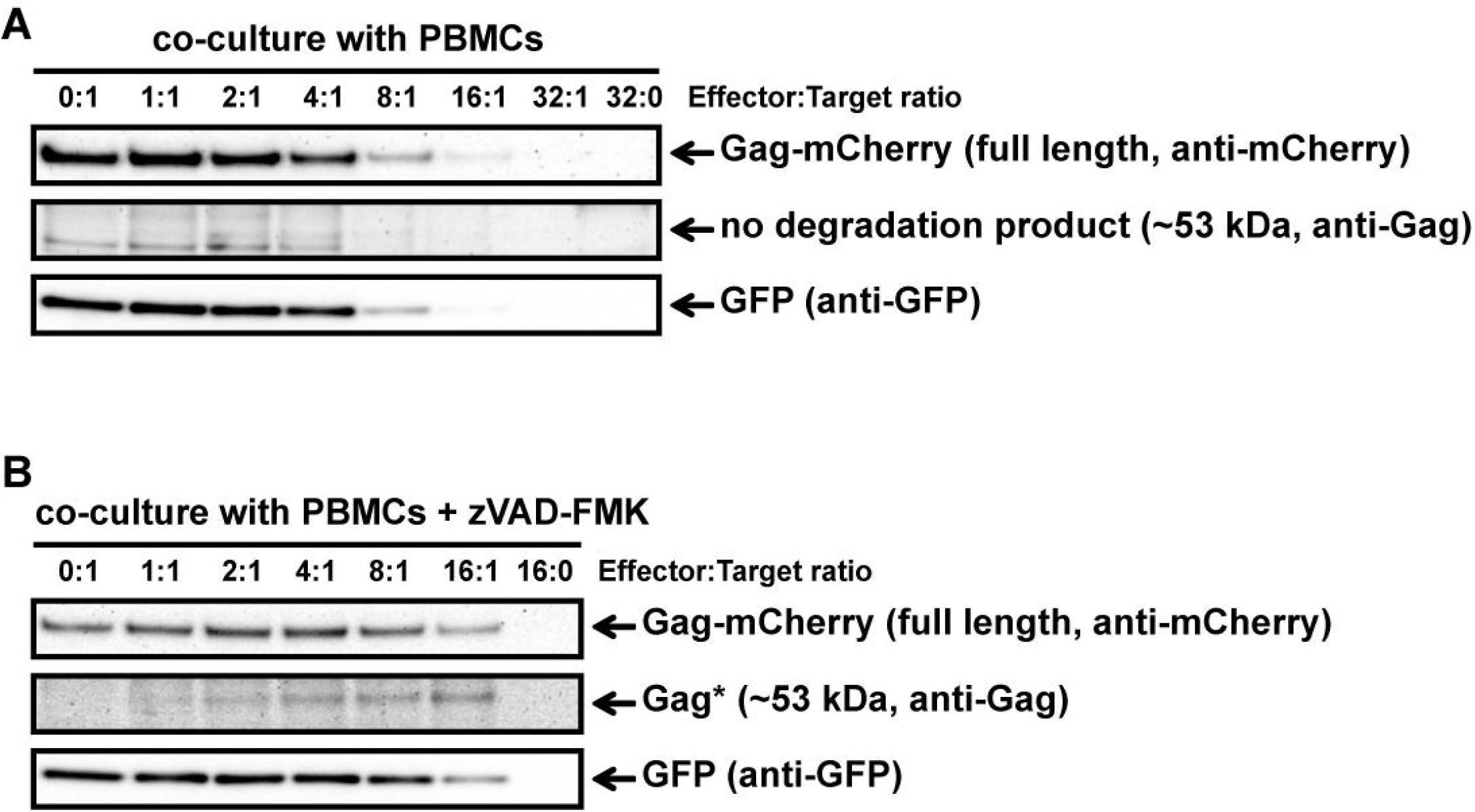
IL-2-activated PBMCs target Gag_PYKEi_ in living target cells. **(A)** HeLa cells were transfected with both mCherry-tagged HIV-1C Gag_PYKEi_ and pEGFP-N1 in a 3:1 plasmid ratio and at 24 h post-transfection these cells were challenged with increasing effector:target (E:T) ratios of IL-2-activated PBMCs for 4 h at 37°C. Lysates were subjected to immunoblotting using an anti-mCherry, anti-p55 Gag or anti-GFP antibody. **(B)** HeLa cells were transfected with both mCherry-tagged HIV-1C Gag_PYKEi_ and pEGFP-N1 in a 3:1 plasmid ratio and seeded in the presence of zVAD-FMK (100 µM) or only DMSO. Then, 24 h post-transfection, these cells were challenged with increasing E:T ratios of IL-2-activated PBMCs in the presence of zVAD-FMK (100 µM) or only DMSO for 4 h at 37°C. Lysates were subjected to immunoblotting using an anti-mCherry, anti-p55 Gag or anti-GFP antibody.

To have a more GrM-specific co-culture model, we used the natural killer lymphoma KHYG-1 cell line as effector cells instead of IL-2-activated PBMCs in our co-culture assays. Indeed, KHYG-1 cells express high GrM protein levels but very low GrB protein levels as observed by flow cytometry (data not shown) and immunoblotting (23). In the absence of the pan-caspase inhibitor, the ~53 kDa degradation product was observed in all E:T conditions except for the controls (0:1 and 16:0) (Figure 5A). When we performed a time course co-culture at a low E:T ratio (2:1) in the absence or presence of the pan-caspase inhibitor zVAD-FMK, a clear increase of the degradation product over time was observed in the conditions where zVAD-FMK was present (Figure 5B).

**Figure 5.**
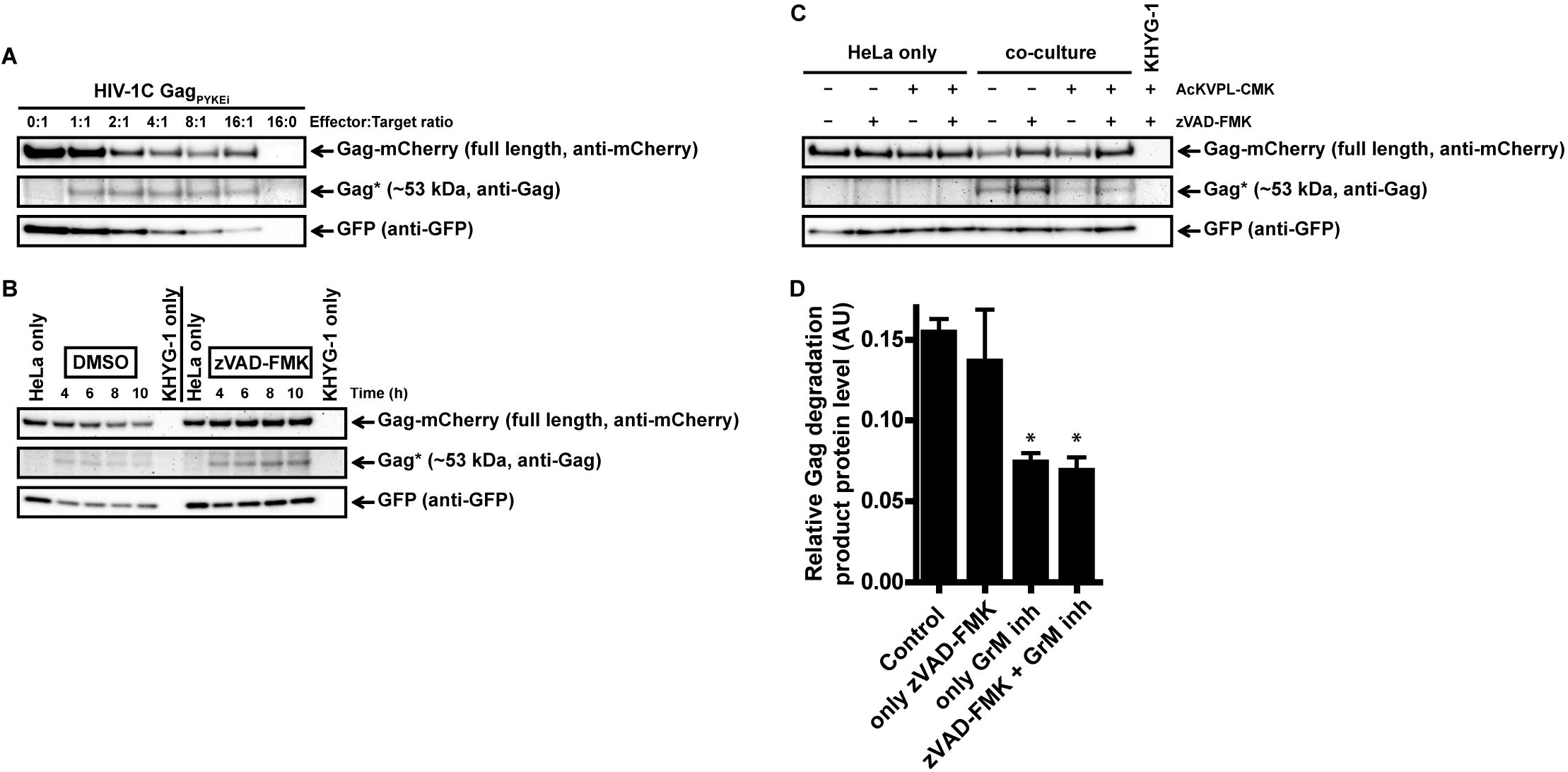
Cytotoxic lymphoma cell line KHYG-1 target Gag_PYKEi_ in living target cells through GrM. **(A)** HeLa cells were transfected with both mCherry-tagged HIV-1C Gag_PYKEi_ and pEGFP-N1 in a 3:1 plasmid ratio and at 24 h post-transfection these cells were challenged with increasing effector:target (E:T) ratios of KHYG-1 cells for 8 h at 37°C. Lysates were subjected to immunoblotting using an anti-mCherry, anti-p55 Gag or anti-GFP antibody. **(B)** HeLa cells were transfected with both mCherry-tagged HIV-1C Gag_PYKEi_ and pEGFP-N1 in a 3:1 plasmid ratio and seeded in the presence of zVAD-FMK (100 µM) or only DMSO. Then, 24 h post-transfection, these cells were challenged with KHYG-1 cells (effector:target ratio of 2:1) in the presence of zVAD-FMK (100 µM) or only DMSO for indicated time points at 37°C. Lysates were subjected to immunoblotting using an anti-mCherry, anti-p55 Gag or anti-GFP antibody. **(C)** Co-cultures were performed as in (b) except KHYG-1 cells were pretreated with GrM inhibitor peptide AcKVPL-CMK (100 µM) or left untreated for 1 h at 37°C before challenging mCherry-tagged Gag_PYKEi_ expressing HeLa cell with KHYG-1 cells for 4 h at 37°C. **(D)** Band intensities of the ~53 kDa Gag degradation product as detected with the anti-p55 Gag antibody in (c) and two additional experiments were quantified, and values were normalized for GFP band intensities. Data points are plotted as mean ±SEM arbitrary units (AU) from triplicate samples. (*p < 0.05 compared to control; ANOVA)

Finally, we performed the KHYG-1 co-culture assay in the presence of GrM-inhibitor peptide AcKVPL-CMK (Figure 5C-D). Degradation of Gag_PYKEi_ was significantly reduced whenever the GrM-inhibitor was present, indicating that this degradation is specifically mediated by GrM. Altogether, these data show that GrM secreted from cytotoxic lymphocytes can cleave Gag in target cells.

### GrM expression within PBMCs of ECs and VPs

Since we showed that GrM can target HIV-1 Gag, this could constitute a mechanism for immunological control of HIV-1. ECs are a group of HIV-1 infected individuals who can control their infection in the absence of antiretroviral therapy, and it has been shown that GrM mRNA could be differentially expressed within PBMCs from ECs compared to viremic progressors (VPs) (18). To assess whether differences in GrM levels might be an underlying mechanism by which EC could control HIV-1, we examined both GrM mRNA transcript levels and protein levels within PBMCs from various patient groups. We collected transcriptomics data on an expanded patient cohort and plotted the expression of GrM transcripts in PBMCs for each individual of HIV-1 negative individuals (HCs), ECs, and VPs. GrM transcript expression was highest in HCs, lowered in ECs and lowest in VPs (Figure 6A). However, these differences were not significantly different. Since perforin is required for the intracellular effects of GrM, we also analyzed the transcript levels of perforin (Supplementary Figure 1). However, no differences in perforin transcript levels were observed among the three patient groups. The same analysis was performed for the transcripts of the other four granzymes (Supplementary Figure 1). Like perforin, GrB transcript levels were similar in all three patient groups. The transcript levels of GrA and GrK of ECs were similar to uninfected individuals and increased in VPs. GrH transcripts were increased in ECs and highest in VPs compared to HCs. Thus, GrM and GrH were the only transcripts differentially expressed in both ECs and VPs compared to the uninfected controls. For all individual granzymes, except for GrK, there was a positive correlation between the granzyme transcript level and perforin transcript levels (Supplementary Figure 2). No correlation was observed between GrM transcript levels with any other individual granzyme transcript levels (Supplementary Figure 3).

**Figure 6.**
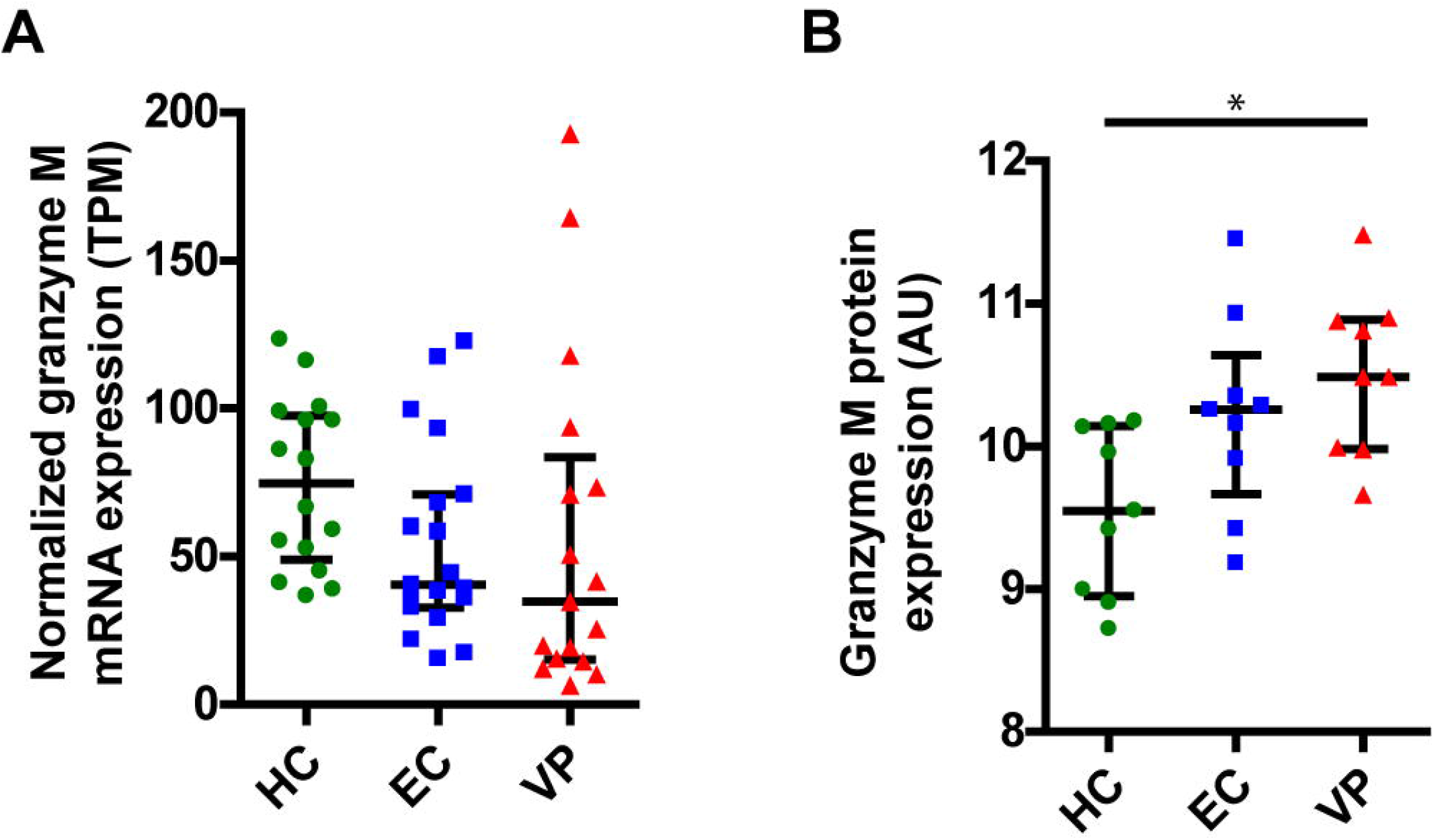
GrM is differentially expressed within PBMCs of individuals from uninfected healthy controls (HC), Elite Controllers (EC) and viral progressors (VP). **(A)** Normalized expression levels (transcripts per million, TPM) of GrM transcripts within PBMCs from each individual are plotted and median with interquartile range is depicted for each patient group. **(B)** GrM protein expression (arbitrary units, AU) within PBMCs from each individual are plotted and median with interquartile range is depicted for each patient group. (*p < 0.05; Kruskal-Wallis)

Next, we examined the protein expression levels of GrM from HCs, ECs, and VPs (Figure 6B). Here, we observed a reversed expression pattern among the patient groups with the lowest GrM protein expression in HCs, increased expression in ECs, and highest expression in VPs. The increased protein expression in VPs was significantly different compared to HCs. Similar to the transcriptomics data, perforin protein levels did not differ between the three patient groups (Supplementary Figure 4). In our proteomics data, GrB and GrK were not detected. Protein levels of GrK were following the same trend as for transcript levels; lowest levels in HC, higher levels in EC, and the highest in VP (Supplementary Figure 4). Thus, there is no clear difference in GrM transcript or protein levels between ECs and other patient groups (HCs and VPs).

## Discussion

Most studies investigating the antiviral response of cytotoxic lymphocytes towards HIV-1-infected cells have focused on GrB-positive CD8^+^ T lymphocytes. Only a few studies have examined the role of other granzymes or perforin, or assessed all five human granzymes in HIV-1-specific CD8^+^ T lymphocytes (11, 24–28). However, these studies only focus on the detection of granzymes within cytotoxic lymphocytes and have not investigated the immunological molecular mechanism by which the granzymes could control HIV-1. Some noncytotoxic antiviral mechanisms employed by granzyme-releasing cytotoxic lymphocytes have been described in the context of other virus infections, such as Herpes simplex virus type 1 (HSV-1), and human cytomegalovirus (HCMV) (8–10). HIV-1 permanently integrates its viral genome into the host cell DNA genome and establish viral latency reservoirs, greatly complicating the possibility to eradicate viruses from the host through cell-mediated cytotoxicity by cytotoxic lymphocytes. However, understanding the role of granzymes in counteracting HIV-1 infection, even through non-cytotoxic mechanisms, and modulating these activities within cytotoxic lymphocytes could be a strategy to control latent or reactivated HIV-1 infection in the absence of antiretroviral drugs or in individuals failing with low-grade viremia during antiretroviral therapy. In our previous study on HIV-1C Gag variants with or without the tetrapeptide insertion PYxE (16), we noticed a potential GrM cleavage site within Gag, and therefore we studied whether and to what extent GrM can cleave Gag in this study.

Here, we indeed show that GrM targets the HIV-1 Gag protein. Even though only a relative small 2 kDa C-terminal part of the P6 late domain is being cleaved off, this exposed domain is important for its interaction with host cell protein ALIX and serves as a binding domain for HIV-1 accessory protein Vpr (16, 29–34). The GrM cleavage site is directly after the ALIX binding motif and cleavage could disturb the interaction of ALIX with Gag and thereby affecting the budding of virions from the plasma membrane (16, 30). The motif for binding Vpr to Gag, LXSLFG, is located after the GrM cleavage site and deletion of this motif completely abolishes the incorporation of Vpr into virions (29, 31–34). It would be of great interest to examine to what extent GrM-mediated cleavage could inhibit virion release or reduce the infectiousness of viral particles.

Although GrM can cleave various variants of Gag, the cleavage efficiency is subtype dependent. Whereas the HIV-1B Gag has a highly conserved amino acid protein sequence near the proline-leucine residues, the HIV-1C Gag shows unique variations with the PYxE tetrapeptide insertions directly before the proline-leucine residues in a subgroup of HIV-1C infected individuals (16). The HIV-1C Gag_PYxEi_ variant seems to be originating from Eastern Africa and is emerging in other countries (16, 35, 36). The PYKE sequence is most likely the result of a recombination event between HIV-1C that lacks this sequence with HIV-2 that naturally contains the PYKE motif. Insertion of the PYKE motif within HIV-1C Gag enhanced its binding capacity to host cell protein ALIX and correlated with increased viral replication and viral fitness (16, 37). On the one hand, PYKE insertion also increases susceptibility towards GrM cleavage. Mutations of the lysine residue into a glutamine (PYQE) or arginine (PYRE) have been observed in different patient cohorts and could be a consequence of immunological pressure (14, 16, 38). Indeed, Gag_PYQEi_ shows reduced susceptibility towards GrM-mediated cleavage. These mutations in the tetrapeptide sequence could be a balanced compromise between increased viral replication and degradation by the immune system. Therefore, immunological pressure through a GrM-mediated cytotoxic lymphocyte response could be an important driving factor on the transmission and evolution of HIV-1C Gag_PYxEi_ variants.

Regulation of granzyme expression is most likely different for each individual granzyme. Indeed, GrM is not upregulated in response to cytokines such as IL-2 or IFNα that upregulates other granzymes (39, 40). Also, expression of individual granzymes vary among virus-specific CD8^+^ T lymphocytes. Human cytomegalovirus (HCMV)-specific CD8^+^ T lymphocytes appear to express higher protein levels of GrM compared to Epstein-Barr Virus-or influenza-specific CD8^+^ T lymphocytes (8). For HIV-1, we observed a decrease in GrM transcript levels in ECs and VPs compared to non-infected individuals, but increased GrM protein levels. The inverse pattern in transcript and protein levels could perhaps indicate a difference in protein turnover and/or release of GrM in cytotoxic lymphocytes. Variations in granzyme levels could also differ in T lymphocytes of each individual depending on the epitope-specificity and/or HLA-specificity of the T lymphocytes that varies from one individual to another. We do see a wide distribution in GrM transcript levels in VPs, although this could be highly influenced by increased immune activation that occurs in VPs.

We also speculated that differences in GrM levels could be an underlying mechanism in ECs to control their infection. Indeed, in a previous study, GrM transcript levels were significantly different between ECs and VPs, but this study had only 6 VPs included in the analysis (18). Here, we expanded the cohort to include 17 VPs. Although there are minor differences in expression of GrM transcripts and protein levels within ECs compared to HCs and VPs, these differences were not significantly different. Of note, we also did not observe any difference in perforin expression between EC and VP in the total PBMC population, although it has been shown that HIV-1-specific CD8^+^ T cells from ECs does express more perforin upon stimulation with HIV-1 peptide pools (28). Thus, our analysis on whole PBMCs might be not sufficient to detect relevant changes occurring in HIV-1-specific CD8^+^ T cells. Also, our limited cohort includes individuals infected by is a mixture of subtypes, including A1, B and C (only wild type) (18). It would be worthwhile to examine whether there is a correlation between the subtype you are infected with, including subtype C with or without the tetrapeptide insertion, and GrM levels within their cytotoxic lymphocytes.

Understanding how the host’s immune response could target HIV-1 through GrM exposes weaknesses of HIV-1 infection that could be exploited therapeutically. Although chemical intervention strategies could be designed that mimic the activity of GrM towards the viral Gag protein, immunotherapies exploiting GrM could be a promising alternative approach. HIV-1-specific CD8^+^ T cells could be genetically modified to increase GrM protein expression and expanded *ex vivo*. However, *ex vivo* or expanded HIV-1-specific CD8^+^ T cells from HIV-1 ECs that are cultured with HIV-1-infected CD4^+^ T cells reduced degranulation activity for all granzyme-positive CD8^+^ T cells (11). This indicates that merely increasing GrM levels in HIV-1-specific CD8^+^ T cells would be insufficient to boost anti-HIV-1-responses. Either *ex vivo* conditions are lacking additional signals that are present at sites of infection to induce CD8^+^ T cell degranulation or the HIV-1-specific CD8^+^ T cells are showing signs of exhaustion. Therefore, it will be important to study which additional signals are required for efficient degranulation of genetically modified GrM-positive CD8^+^ T cells after they have expanded *ex vivo*. Alternatively, if we could stimulate HIV-1-specific CD8^+^ T cells to specifically increase GrM protein expression *in vivo*, we could directly utilize the patients’ immune response to improve anti-HIV-1 activities.

## Acknowledgments

Mass spectrometric analysis and database search for protein identification and quantification were carried out at the Proteomics Biomedicum core facility, Karolinska Institutet, Stockholm.

