## Supplementary Data for "Cytotoxic lymphocytes target HIV-1 Gag through granzyme M-mediated cleavage"

**Supplementary Figures**

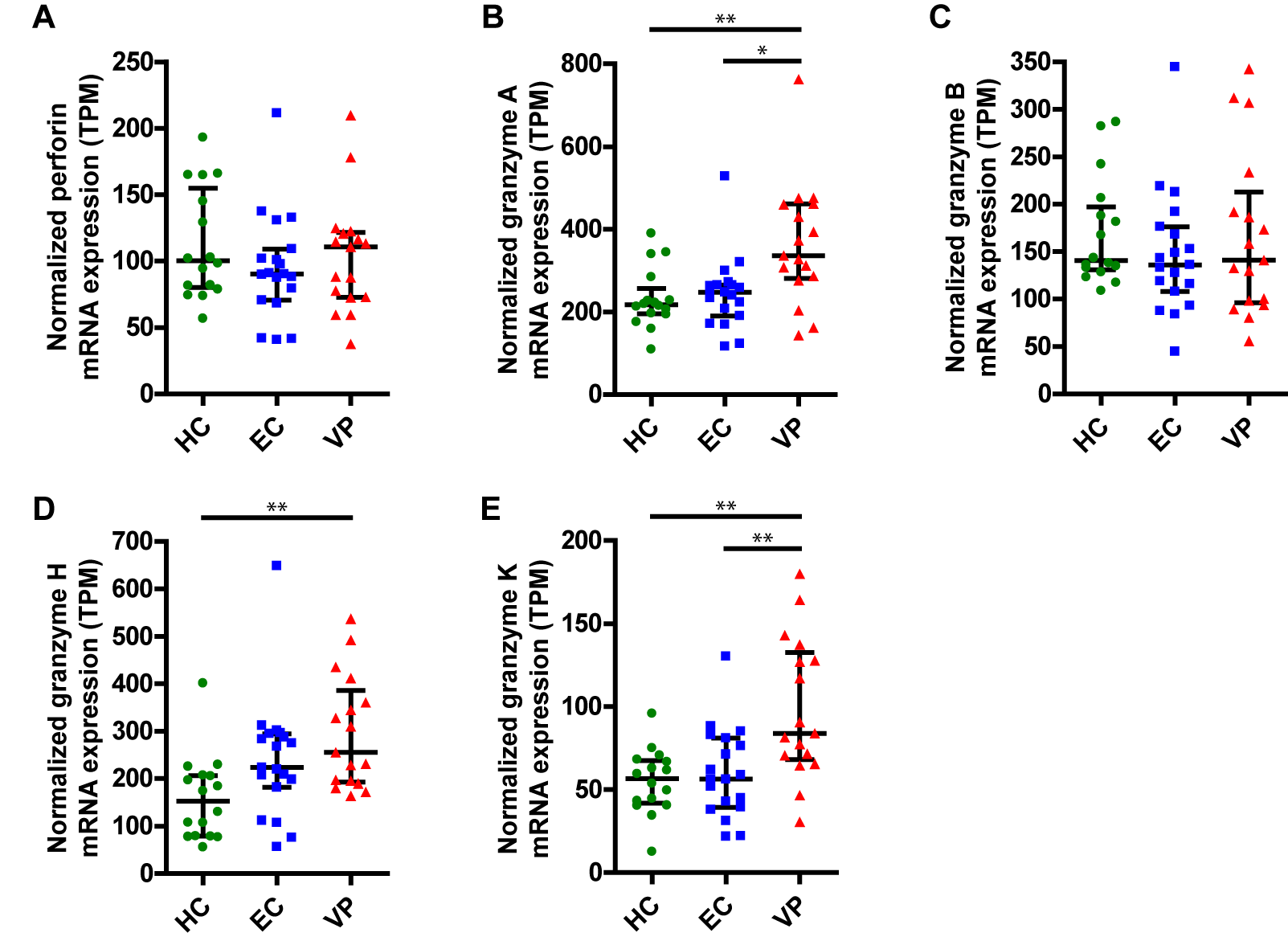

**Supplementary Figure 1. Perforin, GrA, GrB, GrH, and GrK transcript expression within PBMCs of individuals from uninfected healthy controls (HC), Elite Controllers (EC) and viral progressors (VP).** Normalized expression levels (transcripts per million, TPM) of perforin **(A)**, GrA **(B)**, GrB **(C)**, GrH **(D)** and GrK **(E)** transcripts within PBMCs from each individual are plotted and median with interquartile range is depicted for each patient group. (*p < 0.05, **p < 0.01; Kruskal-Wallis)

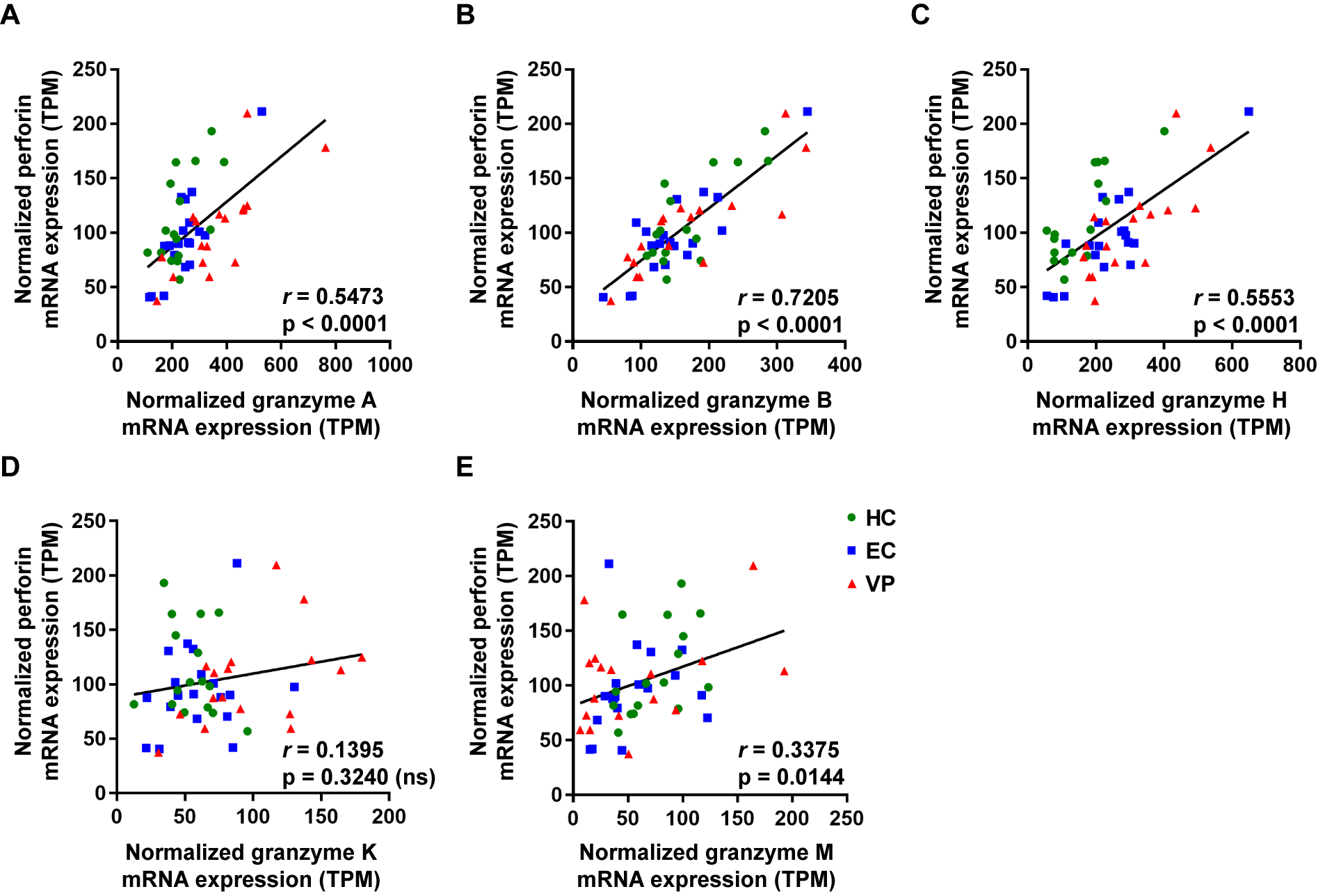

**Supplementary Figure 2. Correlations between perforin transcript levels and each individual granzyme transcript levels.** Normalized transcript levels (transcripts per million, TPM) of perforin against normalized transcript levels of GrA **(A)**, GrB **(B)**, GrH **(C)**, GrK **(D),** and GrM **(E)** from each individual were plotted. Spearman *r* values and p values are depicted within each graph. (ns, not significant)

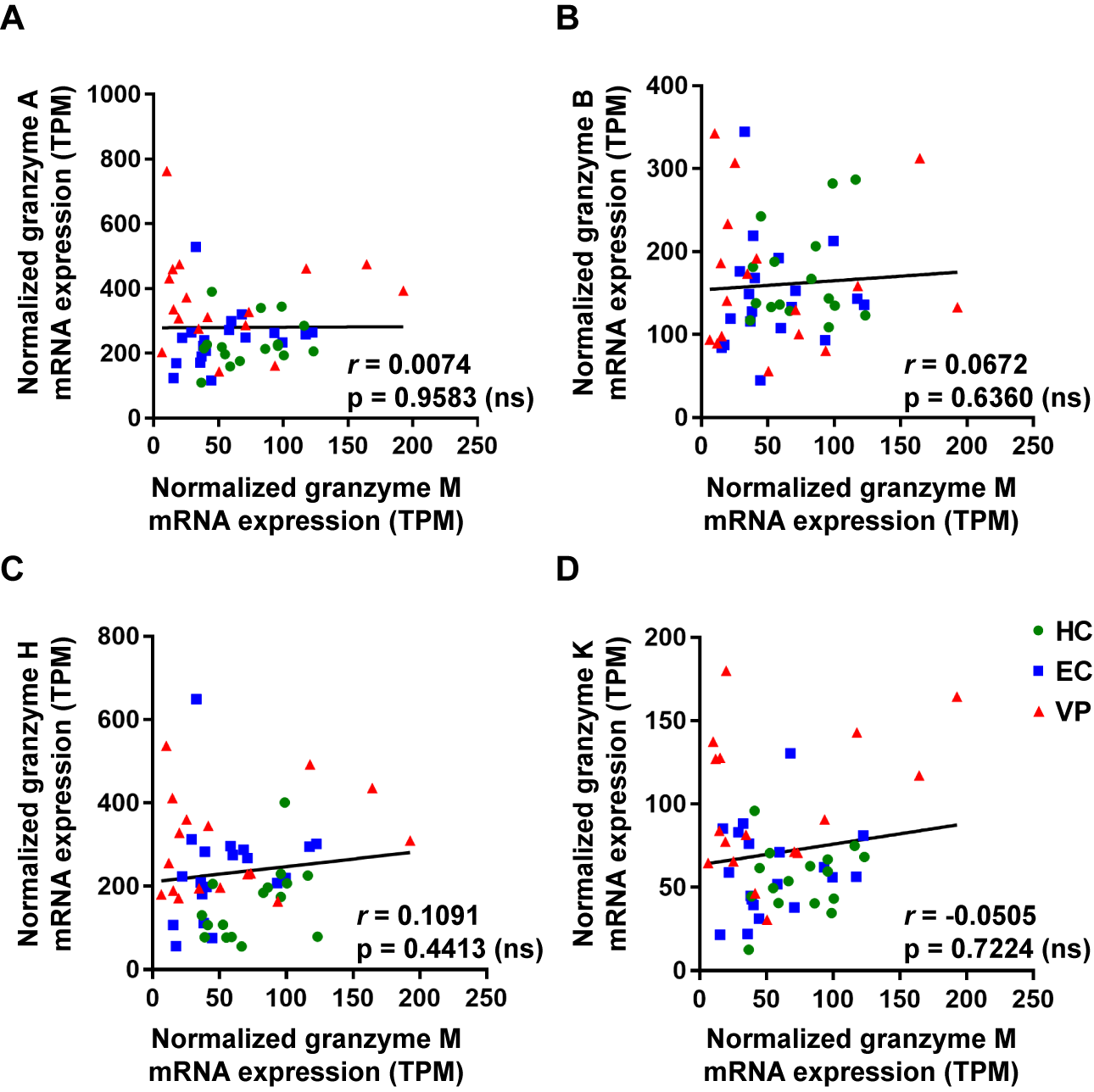

**Supplementary Figure 3. Correlations between GrM transcript levels and each other individual granzyme transcript levels.** Normalized transcript levels (transcripts per million, TPM) of GrM against normalized transcript levels of GrA **(A)**, GrB **(B)**, GrH **(C)**, and GrK **(D)** from each individual were plotted. Spearman *r* values and p values are depicted within each graph. (ns, not significant)

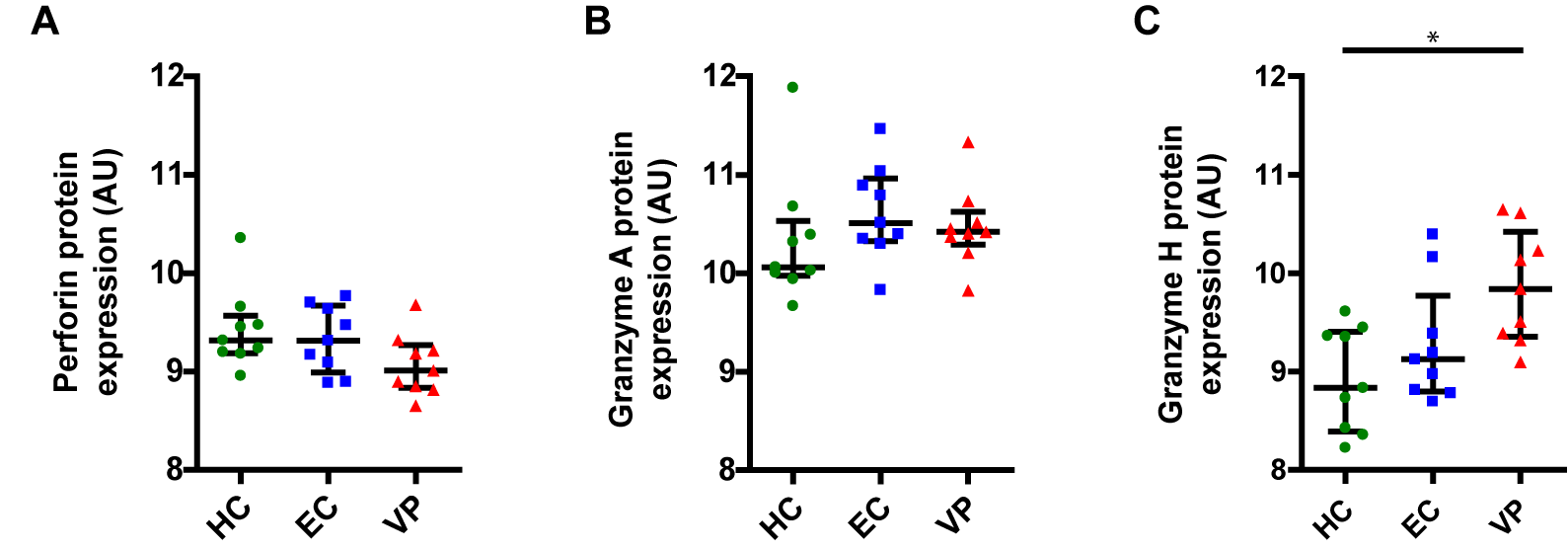

**Supplementary Figure 4. Perforin, GrA and GrH protein expression within PBMCs of individuals from uninfected healthy controls (HC), Elite Controllers (EC) and viral progressors (VP).** Perforin **(A),** GrA **(B)** and GrH **(C)** protein expression (arbitrary units, AU) within PBMCs from each individual are plotted and median with interquartile range is depicted for each patient group. (*p < 0.05; Kruskal-Wallis)
